# The contribution of transposable element insertions to genetic diversity in *Aedes aegypti* populations

**DOI:** 10.1101/2025.10.31.685655

**Authors:** Gabriela Valente-Almeida, Austin Daigle, Daniel R. Matute, Daniel R. Schrider

## Abstract

*Aedes aegypti* is a vector of multiple tropical diseases. The main strategy to control transmission is insecticide-based population control. However, mosquito populations rapidly evolve resistance, possibly enabled by their high levels of genetic diversity. Genome-wide surveys of diversity in *Ae. aegypti* have focused on single nucleotide polymorphisms (SNPs), and although structural variants such as transposable element (TE) insertions have been implicated in insecticide resistance (IR) in *Drosophila*, these have not been thoroughly characterized in *Aedes*. Here, we evaluated the TE content in 122 *Ae. aegypti* genomes from six countries across Africa, North, and South America. We found that TEs contribute substantially to genetic diversity and reflect population structure broadly consistent with that seen in SNPs. Although most TEs insertions are rare, some were observed at higher frequencies, suggesting that a small subset of these may be beneficial. For example, we identified numerous TEs with large frequency differences across populations, consistent with the possibility that these are in haplotypes underlying local adaptation. Specifically, we found three TEs near genes that may be involved in metabolic insecticide resistance: *CYP6P12*, *GSTD11* and *GSTZ1*. In Colombian samples, we also identified a TE insertion that is in negative linkage disequilibrium with several insecticide resistance mutations that form an intermediate-frequency haplotype in the *VGSC* gene region. These results suggest the possibility that, just as TEs have been implicated in adaptation in other animals such as *Drosophila*, they may play an important role in the evolution of resistance to control efforts in *Aedes* and other pests.

## 1. INTRODUCTION

*Aedes aegypti*, commonly known as the yellow fever mosquito, is the primary vector of several tropical diseases, including dengue, Zika, chikungunya, and yellow fever (Viglietta et al. 2021). These diseases impose a significant public health burden, especially in tropical countries (Mitra and Mawson 2017), and recent years have seen numerous epidemics and an increase in case numbers (Lim et al. 2025). For example, 2023 saw a then-record number of dengue cases globally (∼7 million), which was roughly doubled in 2024 (Haider et al. 2025). Due to climate change, adaptation to urban environments, and globalization, it is expected that *Ae. aegypti* will increase its geographic range (Gubler 2011; Kraemer et al. 2019; Ryan et al. 2019). Additionally, mosquitoes are expected to complete their life cycle more rapidly and over longer periods of the year (Iwamura et al. 2020), increasing both their rate of evolution and number of opportunities to contact humans; the latter may lead to higher disease transmission potential. These changes will collectively enhance the vector capacity of these organisms (Iwamura et al., 2020).

Insecticides have been the primary method used to control the population size of mosquitoes. Nevertheless, increasing insecticide resistance (IR) in these populations hinders insecticide efficacy on a long-term basis (Garcia et al. 2018) and consequently threatens the effectiveness of the measures of vector control campaigns. Because of this, identifying the genetic underpinnings of resistance is of major interest to the scientific community, as such information could aid surveillance efforts and interventions to control the mosquitoes’ populations. Numerous studies have identified SNPs that are statistically associated with IR, with several mutations experimentally confirmed as causative (Riveron et al. 2014; Tmimi et al. 2018; Qian et al. 2021). However, much of this work has focused narrowly on SNPs, leaving other forms of genetic variation underexplored. In particular, structural variants (SVs), such as large-scale insertions and deletions, are known to contribute to IR in *Drosophila* (Battlay et al. 2018), with some of these SVs potentially experiencing strong spatially varying selection pressures (Turner et al. 2008; Kolaczkowski et al. 2011; Schrider et al. 2016). Thus, it is possible that SVs make an important contribution to IR in *Aedes* that has yet to be characterized. Among these, transposable element (TE) insertions represent a particularly common class of SVs which can have important fitness consequences (Langley and Charlesworth 1989; Cordaux and Batzer 2009). TEs are mobile DNA sequences capable of excising or replicating themselves and relocating to new positions within the genome, either via copy-and-paste (class I TEs, or retrotransposons) or cut-and-paste (class II TEs, or DNA transposons) mechanisms (reviewed in Wells & Feschotte, 2020). *Aedes aegypti* has a large and highly repetitive genome, with TEs and other repetitive sequences accounting for 65% of its total genomic content (Matthews et al. 2018). Thus, an examination of the extent to which TE insertion polymorphisms (those not yet fixed or eliminated from the population) contribute to genetic diversity in *Ae. aegypti* is warranted.

TEs are known to be associated with deleterious effects as they generate genome instability (Montgomery 1991; Bhat et al. 2022). Moreover, because they move into different locations of the genome, they can be inserted into protein-coding genes, regulatory regions, or other functional sequences (Finnegan 1992), thereby impacting the expression or function of these elements (Feschotte 2008; Lee 2015). In all these scenarios, the effects of these TE insertions would be deleterious in most cases (Hollister and Gaut 2009). Indeed, TEs have been found to be important risk factors for genetic disorders and cancer (Ayarpadikannan and Kim 2014). However, TE insertions can also occasionally have beneficial effects (Feschotte and Pritham 2007; Casacuberta and González 2013). Indeed, the notion that TEs can be associated with insecticide resistance goes back several decades (Wilson 1993; Rostant et al. 2012). For example, some TEs have been shown to confer insecticide resistance in *Drosophila* (Waters 1992; Chung et al. 2007), and insertions within insecticide resistance genes have been observed in other insects, such as *Helicoverpa armigera* (Klai et al. 2020).

Here, we investigate the contribution of TEs to genetic diversity in *Aedes aegypti* and their potential role in IR. Because of the previously mentioned high TE content in *Ae. aegypti*, we might expect a fair amount of TE insertions that are polymorphic in their presence/absence in this species, with some insertions potentially playing an important role in adaptation and the emergence of IR. We analyze whole-genome sequencing data from 122 *Ae. aegypti* specimens collected from six countries across three continents: Brazil, Colombia, Gabon, Senegal, the USA, and Kenya. These population samples have distinct demographic histories (Kent et al. 2025) and have been exposed to different environmental pressures, including unique histories of insecticide use (Lima et al. 2011; McGregor and Connelly 2020; Granada et al. 2021; Sene et al. 2021), allowing us to examine the potential impact of TEs on evolution and recent adaptation in mosquitoes from different locations and selective environments. We found evidence that TEs make a large contribution to genetic diversity within and between geographic regions in *Ae. aegypti*, with hundreds of thousands of TE insertion polymorphisms being present in the six population samples examined here alone. Our results also suggest that while most TE insertion polymorphisms are probably neutral or deleterious, some TE insertions may play an important role in adaptation, including several candidates found in or near IR genes. These findings underscore the urgent need to further characterize the functional and evolutionary impact of polymorphic TE insertions and other SVs in *Ae. aegypti* and other pest species.

## 2. METHODS

### 2.1. Dataset

To describe the TE composition of different *Aedes aegypti* populations, we examined 122 genomes from six different countries (Lee et al. 2019; Rose et al. 2020) (Figure 1): Brazil (18 specimens), Colombia (24 specimens), USA (28 specimens), Senegal (20 specimens), Gabon (13 specimens), and Kenya (19 specimens). Specific locations can be found in Supplementary Table 1.

**Figure 1:**
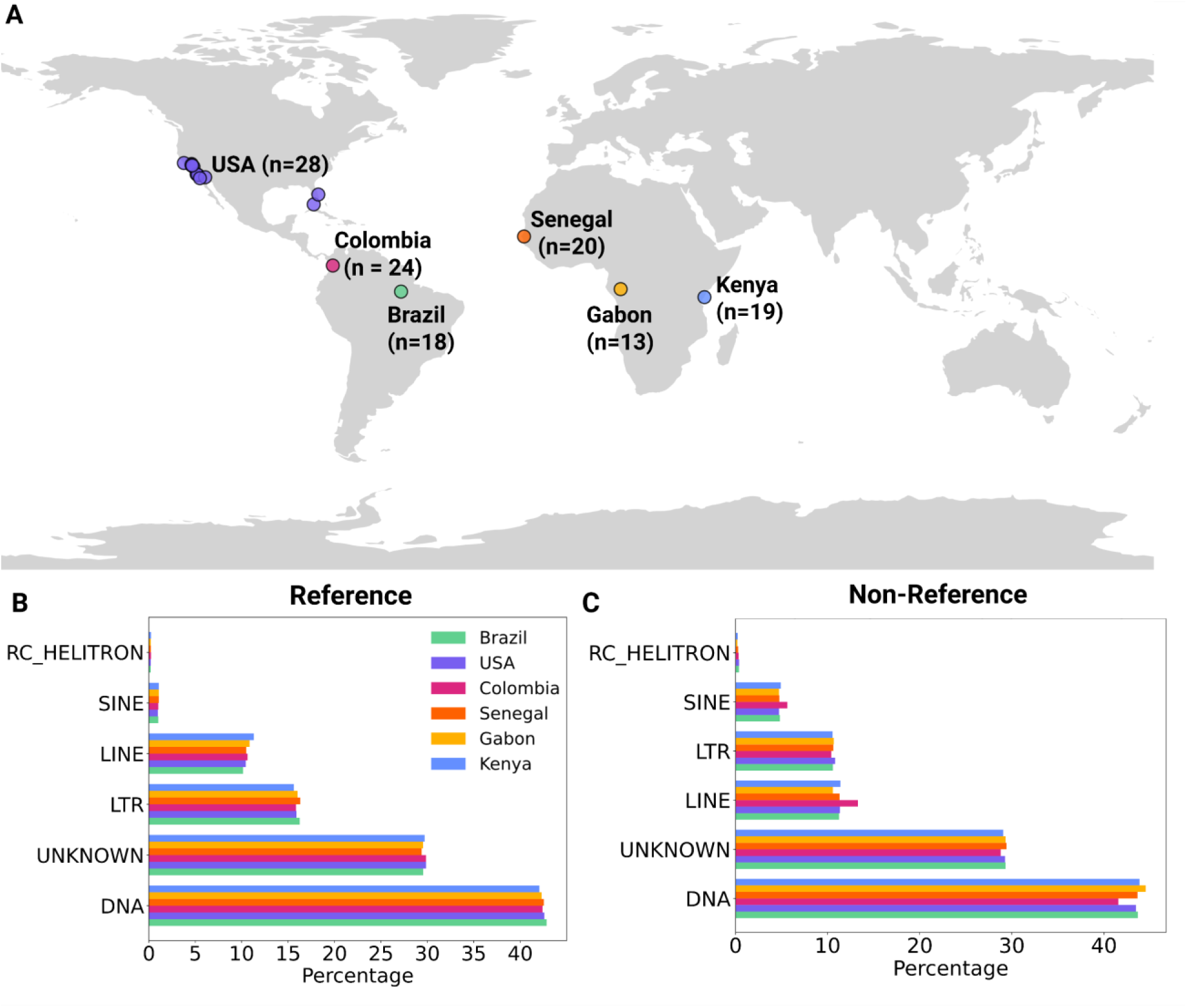
Content of transposable elements in each population of *Aedes aegypti*. A) Map showing the geographic locations of the six population samples used in this study. B) Percentage of reference insertion polymorphisms belonging to each major TE class within each country. C) Percentage of non-reference insertion polymorphisms belonging to each major TE class within each country.

### 2.2. Transposable Element identification - McClintock pipeline

To detect TE insertions, we used the McClintock v.2.0.0 (Nelson et al. 2017) pipeline, which provides a streamlined and user-friendly interface enabling the execution of various methods designed to identify transposable elements from short-read data. These methods for identifying TEs typically classify them into two categories: reference TEs, which are present in the reference genome, and non-reference TEs, which represent insertions absent from the reference. Within this pipeline, we elected to use the methods TEMP2 (Yu et al. 2021) and RetroSeq (Keane et al. 2013).

The reads for each specimen in FASTQ format were retrieved from the NCBI SRA (Supplementary Table 1). Other than the FASTQs, the McClintock pipeline requires a reference genome and a set of consensus TE sequences. We used the AaegL5 reference genome (Matthews et al. 2018), and the TE consensus sequence file from Petersen et al. (2019), which can be found on our GitHub repository (github.com/gabivalentela/TEs_Aedes_aegypti). The consensus sequence file was built using automated annotation methods: RepeatModeler (Smit and Hubley 2008) was used to generate candidate consensus TE sequences, which were then classified using RepBase (Jurka et al. 2005) and further validated with BLAST to confirm their identity as transposable elements. Sequences labeled as “unknown” were those not directly annotated but still showed matches to TE-related proteins such as transposase or reverse transcriptase (Petersen et al. 2019). We configured the McClintock pipeline (Nelson et al. 2017) to automatically annotate the positions of reference TEs using RepeatMasker (Smit et al. 2013). We slightly modified the pipeline so that we could allocate enough RAM to Picard (by adding the -Xmx8g flag to allocate 8 GB when running Picard’s MarkDuplicates).

Non-reference TEs are more challenging to detect than reference TEs because for the latter the location of the possible variant is already known. Previous results showed that TEMP2 (Yu et al. 2021) performs well with *Drosophila* data (Daigle et al. 2025), which initially led us to select it for reference and non-reference TE detection. However, TEMP2 (Yu et al. 2021) includes a filtering step that removes non-reference TEs located in genomic positions that already contain reference TE insertions of the same family, as these may be false positives generated by alignment errors. This resulted in the identification of very few non-reference TEs in our dataset (the mean number of detected non-reference TEs per genome was only 36.52 when using TEMP2). Given that RetroSeq (Keane et al. 2013) also showed good performance with *Drosophila* data (Daigle et al. 2025), we opted to use it for identifying non-reference TEs, while using TEMP2 (Yu et al. 2021) to detect reference TEs.

#### 2.2.1. Detecting reference transposable elements with TEMP2 and subsequent post-processing

TEMP2 detects reference TEs by examining TE insertions that are given as a reference set—in our case, McClintock identified these by running RepeatMasker on the reference genome—and evaluating evidence for the absence of each of these reference TEs across the samples. To determine whether a TE insertion is absent, TEMP2 first selects read pairs whose mate distances are greater than expected based on the average insert size. It then checks whether these read pairs span annotated TEs in the reference genome. To reduce false positives, the observed distance between the read pairs must roughly match the expected insert size after accounting for the excised TE, and the reads must map uniquely to the reference genome. Read pairs supporting the same event are then clustered together, with each cluster representing a putative TE presence/absence polymorphism (Zhuang et al. 2014; Yu et al. 2021).

Our preliminary genotyping results indicated that when multiple TEs of the same type were found near one another, they were often found in the same set of individuals and thus likely represent the same insertion (although we cannot rule out the possibility that they are multiple insertions in strong linkage disequilibrium with one another). In our final genotype set, we therefore considered reference TE insertions to represent the same insertion if they belonged to the same family and type (i.e. reference vs. non-reference) and were located within 1,000 bp of each other in one genome. When two TEs were merged, the start position of the resulting TE insertion was set to the start position of the leftmost TE insertion in the merged pair, and the end position was set to the end of the rightmost TE insertion. This was done repeatedly until no more TEs could be merged for a given individual. However, TEs were not merged across individuals. For example, if one genome contained four TE insertions that were merged, but a second genome had only three of these four TEs, these two genomes were treated as having different TE insertions.

#### 2.2.2. Detecting non-reference TE insertions with RetroSeq and subsequent post-processing

RetroSeq (Keane et al. 2013) detects non-reference TEs in two stages: the discovery phase, where it detects discordant mate pairs and assigns them to a TE class using either annotated TE elements from the reference or alignment with Exonerate to a TE sequence library. Second, the calling phase clusters the TE candidate reads found during the discovery phase based on their mapping location and strand in the reference genome. Finally, predicted TE insertion breakpoints are refined using soft-clipped reads mapping near the insertion site, if any are present.

Because it relies on discordantly mapped mate pairs, a key factor for successful analysis with RetroSeq is having a sufficiently large gap between the two reads at either end of a sequenced insert. For some of the genomes included in our dataset, the insert size was small enough such that the average gap between the reads was very small or nonexistent. We therefore trimmed the original reads so that they would have a max length of 75 bp using FASTP (Chen, 2023), thereby effectively increasing the space between the sequenced reads before running RetroSeq. We previously found this approach to substantially improve performance in genomes where typical inserts had little to no gap between the reads (Daigle et al. 2025). The samples in our dataset had reads that varied in sequence length, so the impact of this trimming step on the effective coverage for TE detection varied from sample to sample and is summarized in Supplementary Table 1.

Some TEs detected by RetroSeq are identified as “hybrid TEs” that show evidence of belonging to two separate TE families. In these cases, we selected the TE family that had stronger read support. To determine which TE family in a “hybrid” insertion call has more support, we used the BAM file to identify all reads mapped within a 2kb window around the hybrid TE’s genomic region. We then matched the read names from this region to those in the RetroSeq discovery file, which contains information about all TE-supporting reads for a genome, including which TE family these reads map to. From these matched entries, we counted how many reads supported each of the two TE families in the hybrid call and chose the family with the highest number of supporting reads as the most likely candidate. If the TEs had the same number of supporting reads, we kept the first TE family name present in the hybrid family label produced by RetroSeq, which follows the same behavior as the default McClintock pipeline (Keane et al. 2013).

To control our false positive rate, we used a breakpoint threshold of 6 (McClintock’s default) when discovering new TEs; this threshold can vary from 1-8 with 8 being the most stringent. After running this initial discovery step to identify candidate TE presence/absence polymorphisms, we then re-ran RetroSeq to perform a more sensitive genotyping step to determine which genomes in our entire data set contained each TE insertion; note again that our genotyping does not distinguish between heterozygous individuals and those homozygous for the presence of the non-reference TE. To avoid an excessively high false negative rate during this genotyping step, we relaxed the breakpoint threshold of 1 (the most lenient one possible) when genotyping TEs already detected in another genome. When doing our genotyping, we considered two non-reference TE insertions detected in different genomes to be the same insertion if they were of the same family and type and within 1,000bp of each other. This merging was performed in the same manner as for reference TEs, except that for non-reference TEs, pairs of TE insertions in different genomes were merged rather than pairs of TEs in the same genome.

### 2.3. Transposable Element polymorphism evaluation and filtering

Previous studies have highlighted the importance of a curation step to ensure the reliability of the transposable elements detected from short reads (David et al. 2024). We therefore incorporated the following curation and evaluation steps before producing our final set of TE insertion calls.

#### 2.3.1. Filtering potential false positives or fixed TE insertions

Our goal was to identify polymorphic TE copies resulting from recent insertions of currently active TE families. However, the TEs identified using McClintock included many reference TE insertions that were fixed across all populations, as well as many reference TEs that were present in all but one individual. Many of the latter insertions may represent truly monomorphic TEs that contained a false positive signature of absence in a single individual, rather than true polymorphisms. In addition, we observed many non-reference TEs present in only a single individual, a category likely strongly enriched for false positives. We therefore applied several filtering steps. First, we retained only TE families that appeared among the non-reference insertions in non-singleton form. Any TE families with only a single non-reference insertion polymorphism found in our dataset were also removed, as we reasoned that these TE families were probably inactive, in which case these TE insertions must be false positives. We also removed reference TEs present in all individuals or in all except one (i.e., the reference TE had to be called absent in 2 or more individuals), as well as non-reference TEs found in only a single individual. Note that these frequency filters were applied to our global population sample as a whole rather than separately to the samples from individual countries. Additionally, non-TE elements such as satellites, simple repeats, and buffer regions were excluded from our final set of genotyped TE insertion polymorphisms.

#### 2.3.2. Determining a minimum length cutoff for reference TE insertions

When a TE is small, there is a higher risk of detecting false-positive TE polymorphisms that seem to be spanned by a moderately large insert and therefore incorrectly appear to be absent, due to natural variation in insert sizes. To evaluate whether this issue was present in our dataset, we plotted site frequency spectra (SFS) for three TE size ranges: those smaller than 200 bp, between 200–600 bp, and larger than 600 bp (Supplementary Figure 1). We found that reference TE insertions smaller than 600 bp had a large excess of high-frequency insertions, suggesting that many of these were likely fixed TE insertions that were incorrectly identified as missing in a small number of genomes. To mitigate the impact of such false positives, we chose to retain only TEs that were 600 bp or larger.

### 2.4. Summaries of population structure and genetic diversity

We summarized the level of TE diversity using standard population genetic statistics and analyses (described immediately below) that were modified to account for the difficulty in confidently distinguishing between heterozygous and homozygous TE insertions. Specifically, each analysis was modified to allow for binary “genotypes” for each TE insertion in each diploid genome: present (whether the TE insertion is present in homozygous or heterozygous state), and homozygous absent.

To summarize within-population diversity, we calculated single-population site frequency spectra. To investigate the extent of population structure present in TE insertion genotypes, we performed principal component analysis (PCA) using scikit-learn (Pedregosa et al. 2011) with default parameters, with the TE insertion genotype matrix used as input. We also calculated *F_ST_* in a pairwise way for all the combinations of countries. The equation used to calculate *F_ST_* was:

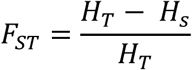

where *H_T_* is the expected heterozygosity when treating the two countries as one population, and *H_s_* is the average expected heterozygosity within each country. To search for TEs potentially involved in population-specific adaptation, we identified TE insertion polymorphisms that vary greatly in frequency between countries. Finally, we plotted joint site frequency spectra for each pair of countries using the scikit-allel package (Miles 2024). When the joint SFS is represented as a 2-dimensional heatmap, a high density along the diagonal indicates similar frequency patterns between countries, suggesting lower divergence, while deviations from the diagonal reflect greater differences in allele frequencies.

We also searched for TEs that may be candidates for contributing to IR by calculating *r*² values between transposable element (TE) insertions and candidate insecticide resistance (IR) nonsynonymous SNPs previously reported in the literature. Specifically, we considered four SNPs—V1016I, I915K, S723T, and V410L—known or suspected to be important for IR in the *VGSC* (Saavedra-Rodriguez et al. 2007; Saavedra-Rodriguez et al. 2008; Harris et al. 2010; Du et al. 2013; Haddi et al. 2017; Saavedra-Rodriguez et al. 2019; Mack et al. 2021) and known to be present in one haplotype in Colombian samples (Love et al. 2023), and one SNP in the *rdl* gene found to be present in intermediate frequencies in US populations and to be associated with resistance to organochlorines (Shotkoski et al. 1994; Feyereisen et al. 2015; Collins et al. 2022; Spadar et al. 2024) we refer to this SNP as A301S according to its position in *Drosophila melanogaster* (following Spadar et al. 2024), although it is located at residue 296 in *Aedes aegypti*. To make the SNP genotypes comparable to the binary TE genotypes, we applied two different encoding strategies. In the first encoding, all heterozygous and homozygous alternate genotypes were considered as “present.” In the second encoding, only homozygous alternate genotypes were considered “present,” while heterozygotes were treated as “absent.” For each encoding, we calculated *r*² values and identified TEs with *r*² > 0.2. We then took the larger of these two values as our *r*² estimate.

## 3. RESULTS

### 3.1. TE insertion polymorphisms are abundant in *Ae. aegypti* populations

We investigated the transposable element content in *Ae. aegypti* genomes sampled from six countries across three continents (Figure 1A). We used the McClintock pipeline to run TEMP2 (Yu et al. 2021) to detect reference transposable element (TE) polymorphisms and RetroSeq (Keane et al. 2013) for non-reference TE polymorphisms (details in Methods). Our initial set contained 1,067,686 reference and 1,326,847 non-reference TE polymorphisms. We sought to limit this set to TE families that seemed to be active, and specifically those TE insertions that appeared to be segregating within the populations. We thus imposed several post-processing steps that included removing fixed TEs, reference TEs only absent in one genome (as that one might also be fixed but not identified due to a genotyping error), and singleton non-reference TEs (see Methods for additional post-processing steps). After imposing these post-processing data quality filters, we obtained a final set of 39,159 reference TEs and 308,659 non-reference TEs. For comparison, our previous analyses of SNPs in this dataset contained 2,810,319 polymorphisms (Love et al. 2023; Kent et al. 2025). Considering the greater size of TE insertions than SNPs, our data suggest that TE insertions are responsible for a much larger total number of polymorphic base pairs than SNPs in *Ae. aegypti*.

The main classes of TEs found in our final set in order of abundance were DNA transposons (labeled “DNA”), TEs that were not previously characterized in *Aedes aegypti* (labeled “Unknown”), retrotransposons that contain long terminal repeats (LTRs), long interspersed nuclear element (LINE) retrotransposons, short interspersed nuclear element (SINE) retrotransposons, and rolling-circle transposable elements (RC Helitrons) (Maringer et al. 2017). The relative contribution of these TE classes was similar for both reference and non-reference TEs, although in non-reference TEs the number of segregating LINEs was higher than LTRs, while the opposite was observed in reference TEs. The fraction of TE insertions derived from each type of TE was similar across all countries (Figure 1B and Figure 1C).

TEs are often associated with genome instability and other factors that typically produce deleterious effects in populations (Finnegan 1992; Feschotte 2008; Lee 2015). However, some TEs have been co-opted by the host to perform important functions and likely experienced positive selection (Gilbert et al. 2021). Our focus here is to infer the possible evolutionary forces acting upon the TE polymorphisms in our dataset. We began by examining the site frequency spectra (SFS) of TE insertions. We found that reference TEs were often observed at intermediate frequencies in the populations (Figure 2A), and that non-reference TEs were predominantly rare (Figure 2B). These patterns are probably influenced by a mixture of ascertainment bias (reference TEs are expected at higher frequencies than non-reference TEs), demographic history, natural selection, and genotyping errors. Thus, the SFS alone does not reveal the selective effect of TE insertions. However, the presence of numerous high-frequency TE insertions does raise the possibility that positive selection may have acted on some of these TEs, either directly or through selection on linked variants. We examine this possibility in sections 3.4 and 3.5.

**Figure 2:**
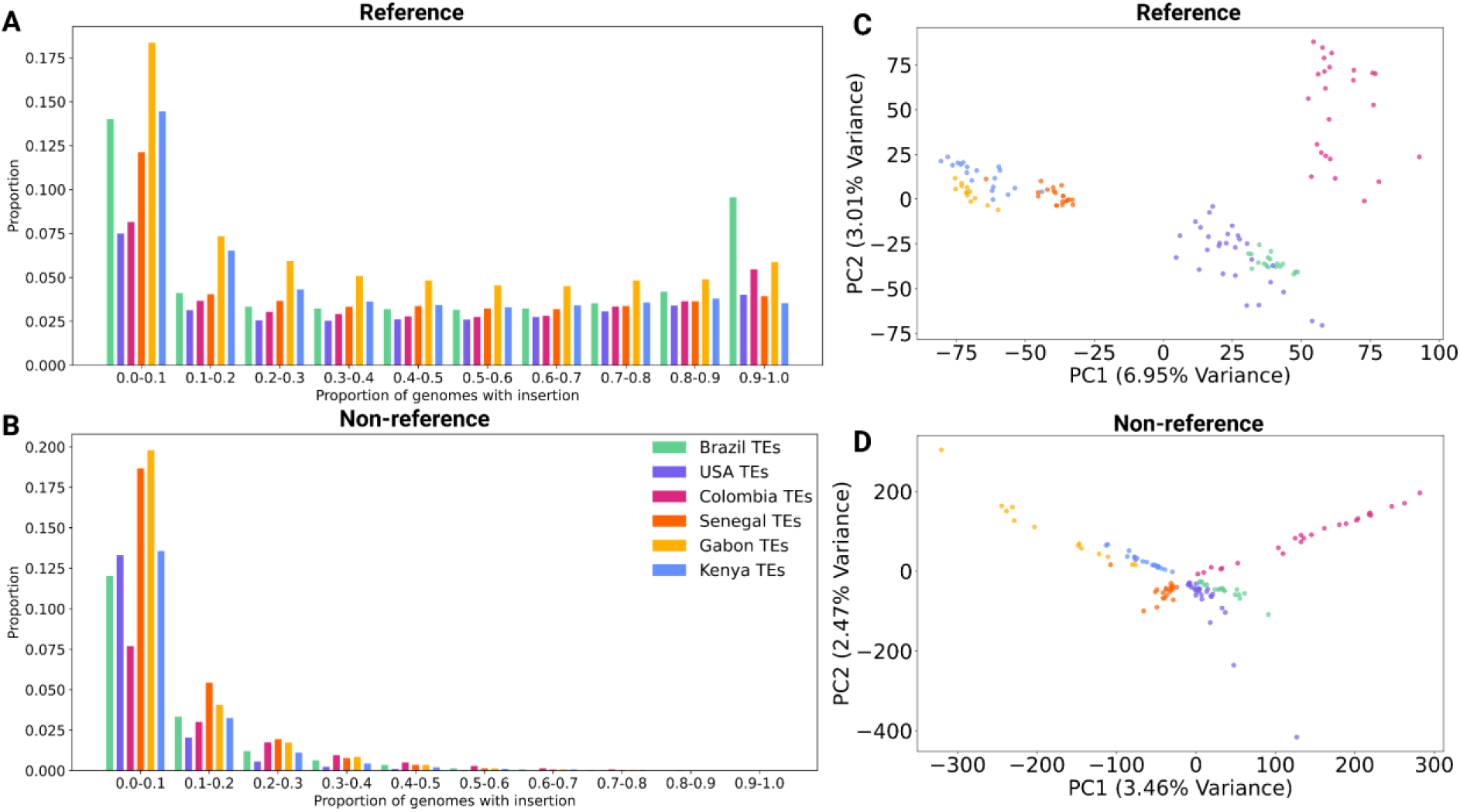
Site frequency spectra (SFS) of transposable element insertion polymorphisms. The SFS was calculated for each country for A) reference TEs and B) non-reference TEs. C) Principal Component Analysis (PCA) of genotypes of reference transposable elements (TEs). D) PCA of genotypes of non-reference TEs.

### 3.2. Transposable element composition reflects population structure in *Aedes aegypti* populations

We sought to assess the extent to which reference and non-reference TEs recapitulate previously observed patterns of population structure in *Ae. aegypti*. First, we used Principal Component Analysis (PCA; see Methods), finding that both reference and non-reference TEs were sufficient to obtain country-specific clusters (Figure 2 C, D). Analyses of SNP data from these same samples had also shown separation by country. It is worth nothing that the top two PCs for TEs explain less of the variation than those obtained for SNPs from the same genomes examined here (Love et al. 2023; Kent et al. 2025). For example, PC1 from Kent et al.’s analysis of SNPs explained 12.35% of the variation (Kent et al. 2025), while our PC1 for reference TEs only explained 6.95% of the variance, and PC1 for non-reference TEs explains only 3.46%. The fact that our reference and non-reference TEs cluster according to geography in a manner that is broadly similar to that previously observed with SNP data suggests that both the SNP and TE PCAs largely reflect the same demographic processes.

Next, we calculated *F_ST_* between each pair of countries for each TE insertion. *F_ST_* values were generally lower between countries on the same continent than between those on different continents (Supplementary Figure 4). The highest *F_ST_* values for reference TEs were between Brazil and Colombia, and Gabon and Colombia (0.14 in both cases). For non-reference TEs and SNPs (data from Love, 2023), Gabon and Colombia had the highest *F_ST_* (0.072 and 0.1 for non-reference TEs and SNPs, respectively). Of the African countries, Senegal showed the least differentiation from the Americas, while of the samples from the Americas, the USA was the least differentiated from the African countries. These results are consistent with Kent et al. (2025), which found that, of the American countries, the USA has the most recent estimated split times with each African country, and that, of the African countries, Senegal has the most recent estimated split times with each country in the Americas. The mean *F_ST_* values for reference and non-reference TEs between African countries (0.04449 and 0.03097, respectively) were very similar to the ones between American countries (0.04505 and 0.03144, respectively), which shows that both continents have a similar level of geographic structure in TE variation. Overall, the pairwise *F_ST_* results corroborate our PCA in that they show that our TE insertion polymorphisms display qualitatively similar patterns of population differentiation seen in SNPs (Love et al. 2023; Kent et al. 2025).

The distribution of *F_ST_* values for a pair of countries can reveal more information than simply examining the mean value—two different pairs could have the same mean genome-wide *F_ST_* but very different distributions. We examined the binned distribution of *F_ST_* values for all combinations between American countries (Supplementary Figure 5A), African countries (Supplementary Figure 5B), between Kenya with all American countries (Supplementary Figure 5C), between Gabon with all American countries (Supplementary Figure 5D), and between Senegal with all American countries (Supplementary Figure 5E). The comparison of African countries revealed some TEs with higher *F_ST_* values (0.8-0.9) while in the Americas, only SNPs were found in that bin. All other combinations between continents showed some TEs within the highest bin (0.9-1.0). Thus, these results suggest that while most TE insertions are not highly differentiated, a minority might show unusual differences in allele frequencies between locations, a possibility we examine in section 3.4.

Next, to examine a richer representation of the degree of differentiation between countries, we calculated the joint site frequency spectrum (joint SFS), which captures the distribution of allele frequencies in two populations simultaneously. The joint SFS for reference TEs (Supplementary Figure 6) and non-reference TEs (Supplementary Figure 7) are broadly consistent with the *F_ST_* values shown in Supplementary Figure 4. Countries within the same continent, such as Brazil and the USA, appear less differentiated, while the joint SFS for intercontinental pairs, such as Kenya and Colombia, displayed more differentiation dispersion away from the diagonal. Among all comparisons, Colombia appeared to be the most divergent from the African countries, consistent with our PCA.

Even after our filtering steps, we observed a large number of reference TEs that are at high frequency in both countries in a given joint SFS. These insertions could be a combination of previously fixed TEs that were recently lost in a small number of samples (e.g., a low-frequency excision of a copy), and truly fixed TEs that were incorrectly genotyped as absent in two or more individuals (thereby surviving our filtering step that removed reference TEs present in all but one individual). We also note that for some pairs of countries, there were a sizable number of TEs at high frequency in one population but low frequency in the other (e.g., Gabon vs Brazil), a point we revisit further below.

### 3.3. Some Transposable Elements Overlap Exons, with Multiple Insertions Detected in Single Exons

To identify TEs that might have deleterious fitness effects, we searched for TE insertions whose estimated breakpoint locations overlapped annotated exons. We found that ∼0.4% and ∼4% of reference and non-reference TEs overlapped with exonic regions, respectively (Supplementary Figure 8 A, C). We further examined the mean frequency of TEs that overlapped exons compared to those that did not, across all populations. Surprisingly, non-reference TEs located within exonic regions had slightly higher mean frequencies than those outside exons, this was observed in the mean frequency calculated using the whole dataset, but also when evaluating each country independently (mean frequency in the whole dataset was 0.15 for non-reference TEs overlapping exons vs. 0.12 for non-exonic non-reference TE insertions). The slightly elevated frequency of exon-overlapping insertions was also observed for reference TEs in Senegal (mean frequency of 0.53 for exonic TEs vs. 0.50 for non-exonic TEs; Supplementary Figure 8B and 8D), but it was not observed in other countries for reference TEs. This unexpected result does not appear to be caused by a bias against detecting and accurately genotyping TEs in repetitive regions (which are less prevalent in exons), as TEs found in exons were found at slightly higher frequencies than TE insertions in non-exonic regions that were not annotated as repetitive (especially for non-reference TEs). Repetitive regions were excluded based on the file containing the annotated repetitive sequences created by Love et al (2023). Although these results may suggest that genotyping accuracy is lower in non-exonic regions even outside of annotated repeats, our results may suggest that the small fraction of TEs within exons that have reached high enough frequency to be detected in our dataset may be no more deleterious than the typical non-exonic TE insertion on average.

### 3.4. Some TE Insertions Differ Widely in Frequency Across Geographic Regions

To explore whether some TE insertions might be subject to spatially varying selection across the geographic regions studied, we examined TE insertions exhibiting the largest frequency differences between countries. Specifically, for each pair of countries, we identified the TEs with the ten highest absolute frequency differences. In total, 126 reference TEs and 102 non-reference TEs showed high variation in frequency between countries. These numbers are lower than the expected 150 (10 TEs x 15 pairs) because some TEs were highly differentiated in more than one country pair. The TEs’ positions, types, and frequency in each country are found in Supplementary Table 2. The mean absolute frequency difference among the 10 most-differentiated TEs for each pair of countries was 0.97 for reference TEs and 0.87 for non-reference TEs.

To investigate the potential functional impact of these differentiated TEs, we examined TEs located within 10 kb of gene regions (84 out of the 126 reference TEs overlapped genes, and 81 out of the 102 non-reference TEs overlapped genes). We then analyzed the GO terms that these genes are annotated with (Supplementary Table 2 and Supplementary Figure 9). Two of the highly differentiated TEs were located near genes (*GSTZ1* and *GSTD11*) annotated with “glutathione metabolic process” and “glutathione transferase” activity, which are cellular processes that can be involved in IR (Lumjuan et al. 2011). *GST* genes, including *GSTZ1*, have been characterized and associated with pesticide resistance in other species (Pavlidi et al. 2018; Ibrahim et al. 2023). *GSTD11* has a TE insertion of unknown type (see Methods for details) located upstream of the gene (Figure 3A). This TE was present only in Colombia and Brazil (96.3% of individuals in Colombia and 22% individuals in Brazil; Figure 3B), showing it could be exclusive to South America. The second gene is *GSTZ1* (Figure 3C), another glutathione transferase—this gene has been shown to be upregulated in insecticide-resistant *Anopheles coluzzii* mosquitoes (Ibrahim et al. 2023). We identified an insertion of unknown type within the intronic region of this gene. This TE insertion is present in all countries but Colombia (present in 42% in Kenya, 50% in the USA, 54% in Gabon, 90% in Senegal, 55% in Brazil; Figure 3D). In addition, we identified a TE located near the cytochrome P450 gene *CYP6P12* (Figure 3E). *CYP6P12* encodes a cytochrome P450 enzyme; members of this enzyme superfamily are known to be involved in pyrethroid resistance, typically through overexpression, which has been shown to enhance insecticide detoxification in multiple species (Waters 1992; Bergé et al. 1998; Scott 1999; Brandt et al. 2002; Daborn et al. 2002; Daborn et al. 2007; Kasai et al. 2014; Yang et al. 2021). The TE was present in higher frequencies in Senegal, Brazil, and Colombia (90% of individuals in Senegal, 39% in Brazil, 62.5% in Colombia, 0% in Gabon, 0% in the USA, and 5% in Kenya; Figure 3F). These TE insertions could potentially impact the expression levels of genes involved in IR in *Ae. aegypti* and thus warrant further examination.

**Figure 3:**
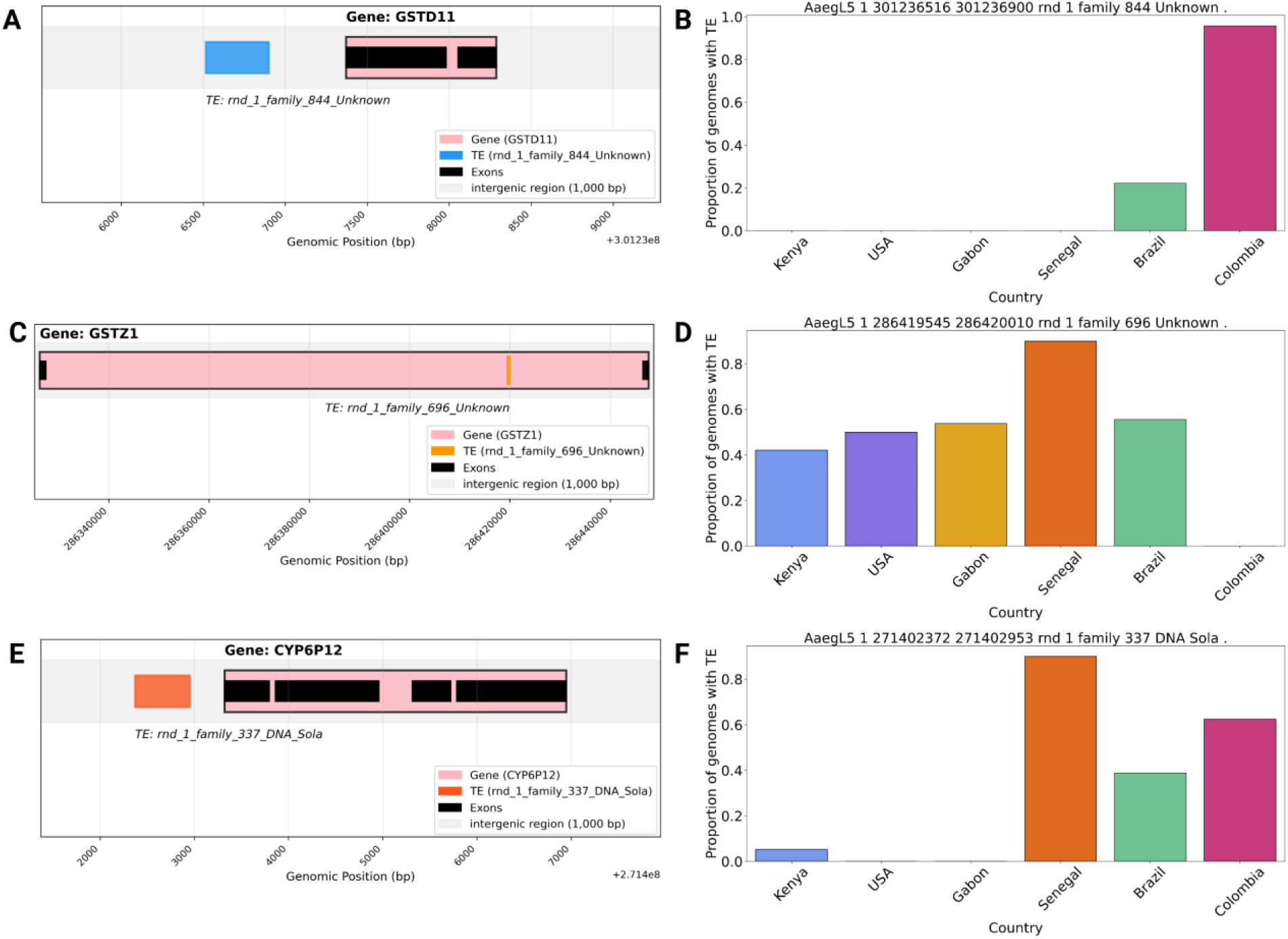
Non-reference TE insertions overlapping gene regions associated with insecticide resistance (IR). A, C, and E) Genomic locations of TE insertions downstream of *GSTD11* (A), within the sequence of the *GSTZ1* (C), and downstream of *CYP6P12*. (E). B, D, and F) Frequencies of these insertions in each of the six countries in our dataset.

### 3.5. Several TE insertions are in linkage disequilibrium with insecticide resistance-related variants in specific populations

Previous studies have shown that TEs can play a role in IR (*e.g.* Daborn et al. 2002; Salces-Ortiz et al. 2020). We therefore explored whether any of the TE polymorphisms in our set were in or near known IR genes. Specifically, we focused on two genes with known resistance-associated variants: *VGSC* (voltage-gated sodium channel; Tognarelli et al. 2025) and *rdl* (resistance to dieldrin locus; Shotkoski et al. 1994; Feyereisen et al. 2015; Collins et al. 2022; Spadar et al. 2024). These genes were selected because they are well characterized in terms of insecticide resistance in *Aedes aegypti* and harbor previously identified nonsynonymous IR variants in the countries represented in our dataset. Other important IR genes (such as, *ACE-1* and *GSTe2*) were not included in this analysis as we did not find any experimentally characterized resistance-associated variants in the literature for these genes that were also present in our dataset. Motivated by our previous observation of strong LD among resistance mutations in *VGSC* in Colombia (see below), we first asked whether any TEs located within 10 kb of these genes are in linkage disequilibrium with variants previously implicated in resistance, as any such insertions could potentially be involved in IR.

The *VGSC* gene is well-known for its role as an insecticide target, and several *VGSC* SNPs have been identified that are strongly associated with IR—particularly to pyrethroids—in *Aedes* and other insects (Clarkson et al. 2021; Diallo et al. 2021; Valmorbida et al. 2022; Spadar et al. 2024). In addition, we previously found that the Colombian sample in our data set contained a resistance haplotype with four known or suspected IR variants in strong LD with each other (Love et al. 2023): V410L (Haddi et al. 2017; Saavedra-Rodriguez et al. 2019), S723T (Saavedra-Rodriguez et al. 2019; Mack et al. 2021), I915K (Mack et al. 2021), and V1016I (Saavedra-Rodriguez et al. 2007; Saavedra-Rodriguez et al. 2008; Harris et al. 2010; Du et al. 2013). Two of these mutations—V1016I (Du et al. 2013; Hirata et al. 2014; Chen et al. 2019) and V410L (Haddi et al. 2017)—have been experimentally shown to confer pyrethroid resistance. We identified one reference TE insertion in strong negative LD with these variants: a DNA CMC Chapev element located in an intron 76,918bp away from V410L (Figure 4A). The *r*² value was of 0.713 with V410L, which we used to tag the entire resistance haplotype. Although our TE genotyping does not distinguish between heterozygotes and homozygotes for presence of the insertion, and LD estimates may be impacted by genotyping error, these results are consistent with the possibility that this CMC Chapev insertion is in strong negative LD with the four aforementioned IR variants. This in turn raises the possibility that the absence of this TE insertion impacts the degree of resistance conferred by the IR haplotype found in Colombia.

**Figure 4:**
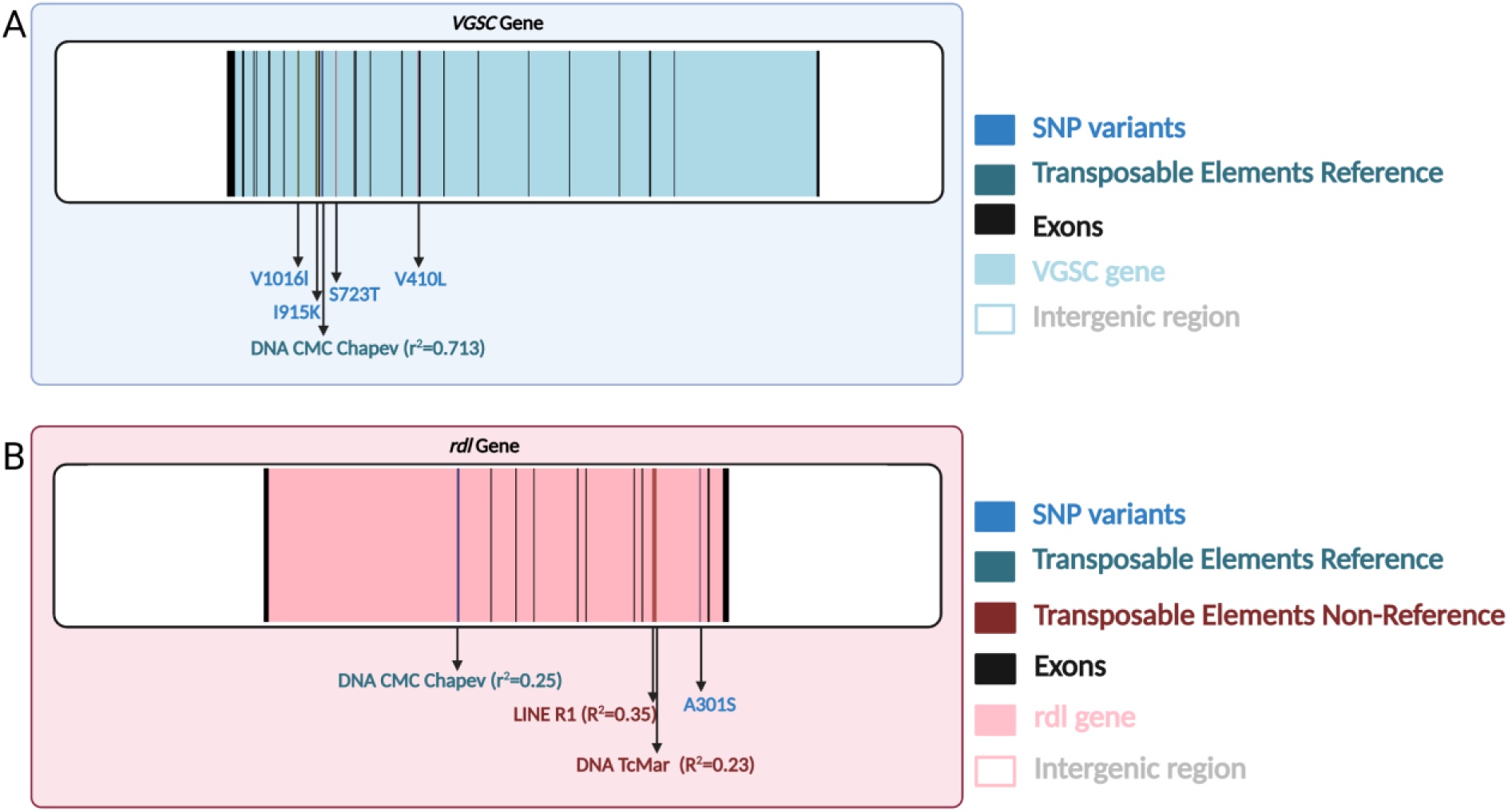
A) TE insertions found to be in linkage disequilibrium (LD) with known insecticide resistance (IR) polymorphisms. A) A reference TE insertion in strong LD with the V410L IR mutation in *VGSC* in the Colombian sample. The locations of other candidate resistance SNPs in LD with V410L in Colombia (V1016I, I915K, and S723T) are also shown. B) TE insertions found to be in LD with the A301S variant in the *rdl* gene in the US population.

Another gene associated with IR is *rdl*, which encodes a subunit of the GABA receptor complex and has been linked to resistance phenotypes (Spadar et al. 2024). One previously characterized variant, A301S, is found at intermediate frequencies in U.S. populations and is associated with resistance to organochlorine insecticides (Spadar et al. 2024). We searched for TEs in LD with this variant in our USA sample, but did not identify any TEs in as strong LD as observed for the TE insertion in *VGSC* described above. However, we did find several TEs in moderate LD with A301S: a DNA CMC Chapev reference TE located in an intron 121,464 bp away from A301S (*r*^2^= 0.25 in the USA sample; Figure 4B), and two non-reference TEs, a LINE R1 (24,087 bp away from the variant; *r*² = 0.35) and a DNA TcMar (22,169 bp away; *r*² = 0.23). Genotype information for all TEs and IR variants examined in this section is provided in Supplementary Tables 3 and 4. These TE insertion polymorphisms in the *VGSC* and *rdl* gene regions could represent important insertions that may be involved in IR; experimental analysis will be required to determine if this is the case.

To search for additional TE insertion polymorphisms that may be involved in IR, we identified high-frequency TEs (present in at least 50% of the individuals in each population sample) located within other genes known to be associated with IR (Supplementary Table 5), as such variants may be enriched for those that were driven to high frequency by positive selection. This analysis is complementary to our examination of highly differentiated TEs above as it identifies insertions that have risen to high frequency in a focal country without regard to its frequency in other countries. Cytochrome P450 genes showed the highest number of high-frequency TE insertions: we found 13 TE insertions within 11 *CYP* genes. Other genes like *VGSC* and *NVY* contained multiple TEs (9 insertions in the *VGSC* gene and 10 insertions in the *NVY* gene). *NVY* is a transcriptional regulator that has been seen to be associated with permethrin resistance (Campbell et al. 2019). This set of TE insertions may be enriched for IR mutations and thus represent candidates for further study.

## 4. DISCUSSION

TEs are key drivers of genome evolution and have been implicated in adaptation (González and Petrov 2009), yet their role in shaping population-level diversity in *Aedes aegypti* and other important vectors remains unexplored (Daron et al. 2025). Here, we show that polymorphic TE insertions contribute significantly to genetic diversity in *Ae. aegypti* population samples collected from six different countries. TEs alone were sufficient to produce country-specific clusters in the PCA, consistent with the patterns obtained using SNPs. While most TE insertion polymorphisms are probably neutral or deleterious, we identified numerous candidate insertions that may be adaptive. These include TEs that varied widely in frequency between countries, some of which were inserted in or near genes that may play a role in insecticide resistance (*GSTZ1*, *GSTD11*, and *CYP6P12*). These findings elucidate the composition of TEs in the genomes of *Aedes aegypti* and their variation among different individuals and countries, and reveal new possible candidate IR mutations for further study. We discuss these results and their implications in more detail below.

The *Aedes aegypti* genome is highly repetitive, with about 65% of the reference genome composed of annotated repetitive elements (Matthews et al. 2018). Indeed, the larger genome size in *Aedes* compared to other mosquito genera like *Anopheles* is largely due to TE-driven expansion (Sinkins 2007).. While previous studies have shown that TE content is highly variable across mosquito species, and that their genomes can be invaded through horizontal transfer (Melo and Wallau 2020), no previous studies have characterized genome-wide variation in TE content within *Aedes aegypti* populations. However, the content of TEs has been previously explored in 120 individuals of *Aedes albopictus* from different populations via transposon display genotyping, revealing the presence of population structure between different geographic locations (Goubert et al. 2017). Similarly, here we found that performing PCA on TE insertion genotypes in *Ae. aegypti* produces clusters corresponding to geographic location (Figure 2 C, D), following a broadly similar pattern as that observed for SNPs (Love et al. 2023; Kent et al. 2025). However, our TE data showed some differences from previous analyses of SNPs. For example, in our PCA, the Colombian sample showed greater separation from other samples from the Americas than previously seen in SNPs (Love et al. 2023; Kent et al. 2025). This may be a result of increased genetic drift in Colombia (which experienced stronger bottlenecks than other samples; (Kent et al. 2025)) and/or recent changes in the activity of certain TE families in this sample, although detection/genotyping error may play a role. It is also possible that geographic differences in infection rates may impact TE content, as viral infection of *Ae. aegypti* mosquitoes has been shown to modulate TE transcript levels, with different viruses are associated with different directions of modulation (Garambois et al. 2024).

We observed many reference TE insertions at intermediate or relatively high frequencies (Figure 2A), a finding that a sizeable fraction of these variants are not strongly deleterious (Arkhipova 2018). This is perhaps expected given the gene-sparse nature of the *Ae. aeypti* genome, as most new TE insertions may be unlikely to disrupt functional elements. Indeed, A recent study examining the distributions of sequence divergence estimates between individual TE copies and the consensus sequence in the *Anopheles gambiae*, *D. melanogaster*, and *Ae. aegypti* reference genomes revealed an apparent skew toward somewhat older TE insertions in *Aedes* in comparison to the other two species (Daron et al. 2024), lending further credence to the possibility that insertions in Aedes are less likely to be quickly removed by purifying selection. We also searched for evidence of adaptive TE insertions, including those potentially involved in IR, as has been observed in *Drosophila* (Daborn et al. 2007; Mateo et al. 2014). We focused on IR because these populations are actively adapting to this strong selective pressure (Valle et al. 2019), and because emerging resistant populations have major implications for human health. TEs have been thought to impact IR through multiple mechanisms, such as altering transcriptional regulation, affecting mRNA stability, modifying splicing patterns, or causing loss of gene function (reviewed in ffrench-Constant 2023). A well-known example is the increased metabolic resistance to insecticides in *Drosophila* due to a TE insertion near the *Cyp6g1* gene, which leads to upregulation of this cytochrome P450 enzyme in the fly’s midgut (Daborn et al. 2002; ffrench-Constant 2023). Out of 182 potentially IR-related genes that we examined, we found reference TE insertions within the gene region of 17 genes and non-reference TE insertions in 19 genes. These insertions therefore represent candidates for further study to elucidate any role in IR.

To identify TE insertion polymorphisms potentially involved in local adaptation, we examined insertions that were highly differentiated in genotype frequency between pairs of countries. These highly differentiated TEs included insertions near or in glutathione-s-transferase (GST) and cytochrome P450 monooxygenases (CYP) genes known to be associated with enhanced insecticide metabolism (Saavedra-Rodriguez et al. 2021). These include *CYP6P12*, which has been shown to be strongly associated with pyrethroid resistance when overexpressed in *Aedes albopictus* (Ishak et al. 2016), and *GSTD11* and *GSTZ1*, these are part of the GST gene family that includes *GSTe2* genes, for which elevated expression is associated with resistance to DDT and pyrethroids (Lumjuan et al. 2011). The insertion in the *GSTD11* gene has only been found in Colombia and Brazil (present in 96% and 22% of genomes, respectively); thus, if this TE insertion plays a role in IR (or is in LD with an IR mutation), this specific resistance mechanism could be restricted to South America. *GSTD11* may be involved in resistance to multiple insecticides, including temephos, a larvicide used worldwide to control tropical diseases (Viana-Medeiros et al. 2018). Temephos resistance has been reported in Mexico, Cuba, Brazil, and Colombia (Braga et al. 2004; Grisales et al. 2013; Davila-Barboza et al. 2024; Piedra et al. 2024) and cytochrome P450 genes have been implicated to temephos resistance in Colombia (Grisales et al. 2013). Our results motivate future studies to determine whether *GSTD11* underlies a novel resistance mechanism in South American *Ae. aegypti* populations, perhaps mediated by the TE insertion we identified here. We note that although we concentrated on genes previously associated with IR, other adaptive processes may also be at play, and indeed some highly differentiated TEs were found near genes that overlapped with genes potentially unrelated to IR (Supplementary Figure 9). We also found one TE insertion in strong LD with four previously described insecticide variants in the *VGSC* gene, V1016I, I915K, S723T, and V410L (Saavedra-Rodriguez et al. 2007; Saavedra-Rodriguez et al. 2008; Harris et al. 2010; Du et al. 2013; Haddi et al. 2017; Saavedra-Rodriguez et al. 2019; Mack et al. 2021). *VGSC*, the voltage-gated sodium channel, is one of the main targets of multiple insecticides, including pyrethroids. Pyrethroids kill insects by extending sodium channel activation, which results in the inhibition of action potential propagation in neurons, causing paralysis and death (Davies et al. 2007). Mutations in the *VGSC* gene can confer knockdown resistance (Du et al. 2016). We previously reported that the four IR mutations noted above are in strong LD with each other in Colombia, along with evidence of a recent, and potentially ongoing, sweep of a haplotype carrying the four IR mutations noted above (Love et al. 2023). Our findings suggest that this haplotype may also be identified by the absence of the CMC Chapev insertion. This absence allele of this TE insertion could thus have an effect on IR, or it may be merely hitchhiking with the true targets of selection, a distinction that could require experimental genetic studies to resolve.

We encountered several difficulties in using short read-based callers to identify segregating TE insertions, which was expected given the highly repetitive nature of the *Ae. aegypti* genome (Matthews et al. 2018). However, we found that careful curation was able to somewhat mitigate these issues. To prevent false negative reference TEs (TEs present in some genomes but called as absent) from misclassifying fixed TE insertions as polymorphic, we filtered out small reference TE insertions. For non-reference TEs, we focused only on TE families that showed strong evidence of being active, and for both categories of TE, we took additional filtering steps to enrich our set of variants for true TE polymorphic insertions (Methods). After performing this filtering, we found that, in addition to presenting site frequency spectra that appear less biased, our TEs broadly reflected patterns of population structure seen in SNP data. Thus, when combined with careful curation, short read-based methods for detecting TE insertions can be used to capture true patterns of diversity in TE content in spite of technological challenges, even in a highly repetitive genome such as *Ae. aegypti.* We therefore argue that this strategy could prove effective in other vector or pest species, and may uncover further TE insertions associated with insecticide resistance and other important phenotypes.

The results of our analysis of TE insertion polymorphism in *Aedes aegypti* genomes from six countries indicates that TEs merit greater attention in the study of insecticide resistance, where research has traditionally focused on SNPs. More broadly, structural variants, including TEs, could be key contributors to adaptation in vector populations, as has been observed in other insects like *Drosophila* (Demuth and Hahn 2009; Casacuberta and González 2013; Battlay et al. 2018). Future work will benefit from more samples from other regions in the world, more sequenced individuals, and more species—given the evidence of horizontal transfer of TEs in mosquitoes (Melo and Wallau 2020), the latter may also prove useful for identifying the sources of TEs in vector species. Using long-read sequencing of population samples will be critical for generating more comprehensive catalogs of TE and structural variant diversity. This in turn will enable finer-scale inference of the role of TEs in the evolution of resistance to control efforts, rapidly changing environments, and other selective pressures encountered by *Aedes* and other important vector species.

## DATA AVAILABILITY

All scripts used for the analyses, together with the reference annotation files and the consensus TE file used in this study, are available at https://github.com/gabivalentela/TEs_Aedes_aegypti.

