## Supplementary Figures for "The contribution of transposable element insertions to genetic diversity in *Aedes aegypti* populations"

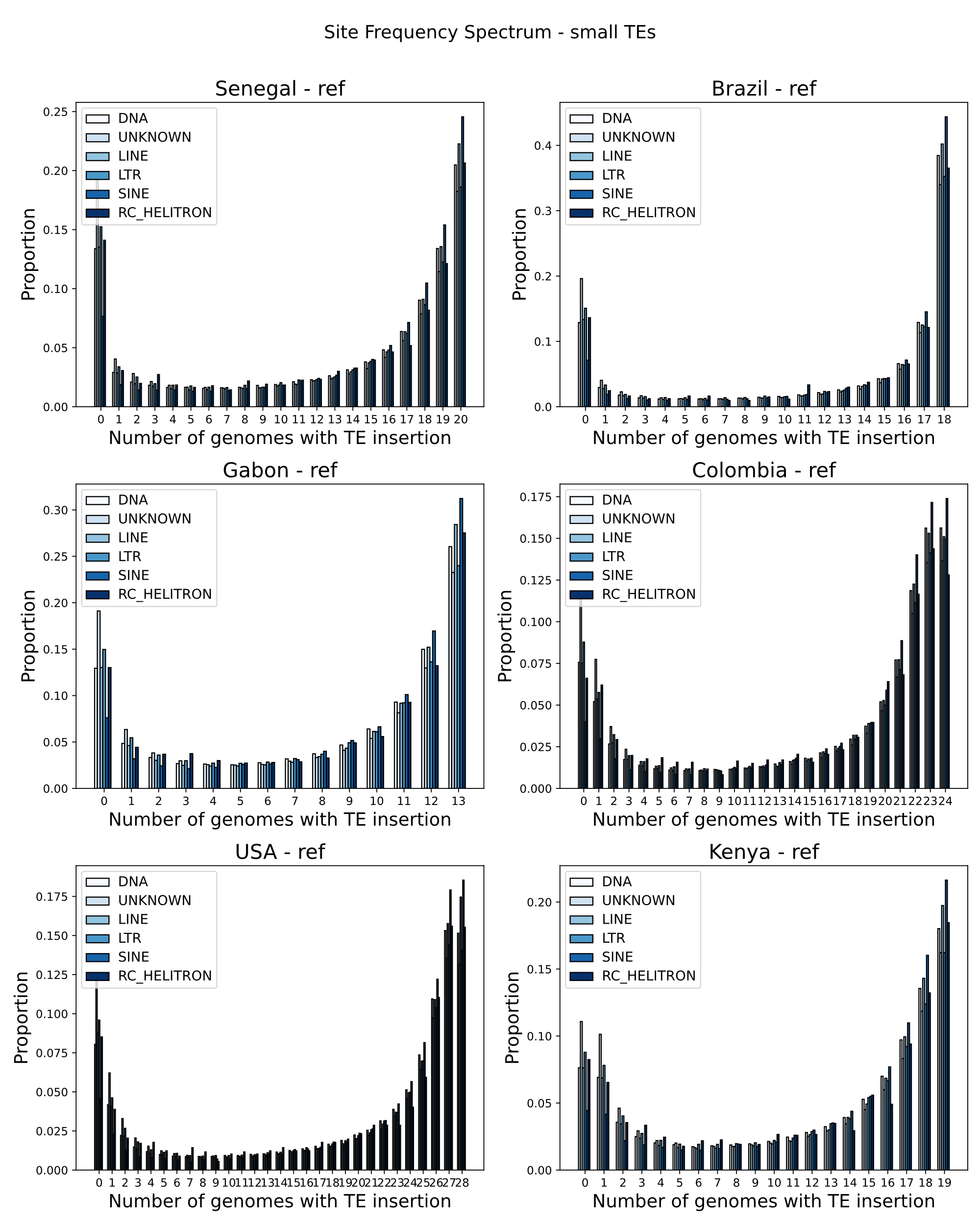

**Supplementary Figure 1:** Site Frequency spectrum of reference TEs based on the length. Smaller than 200bp.

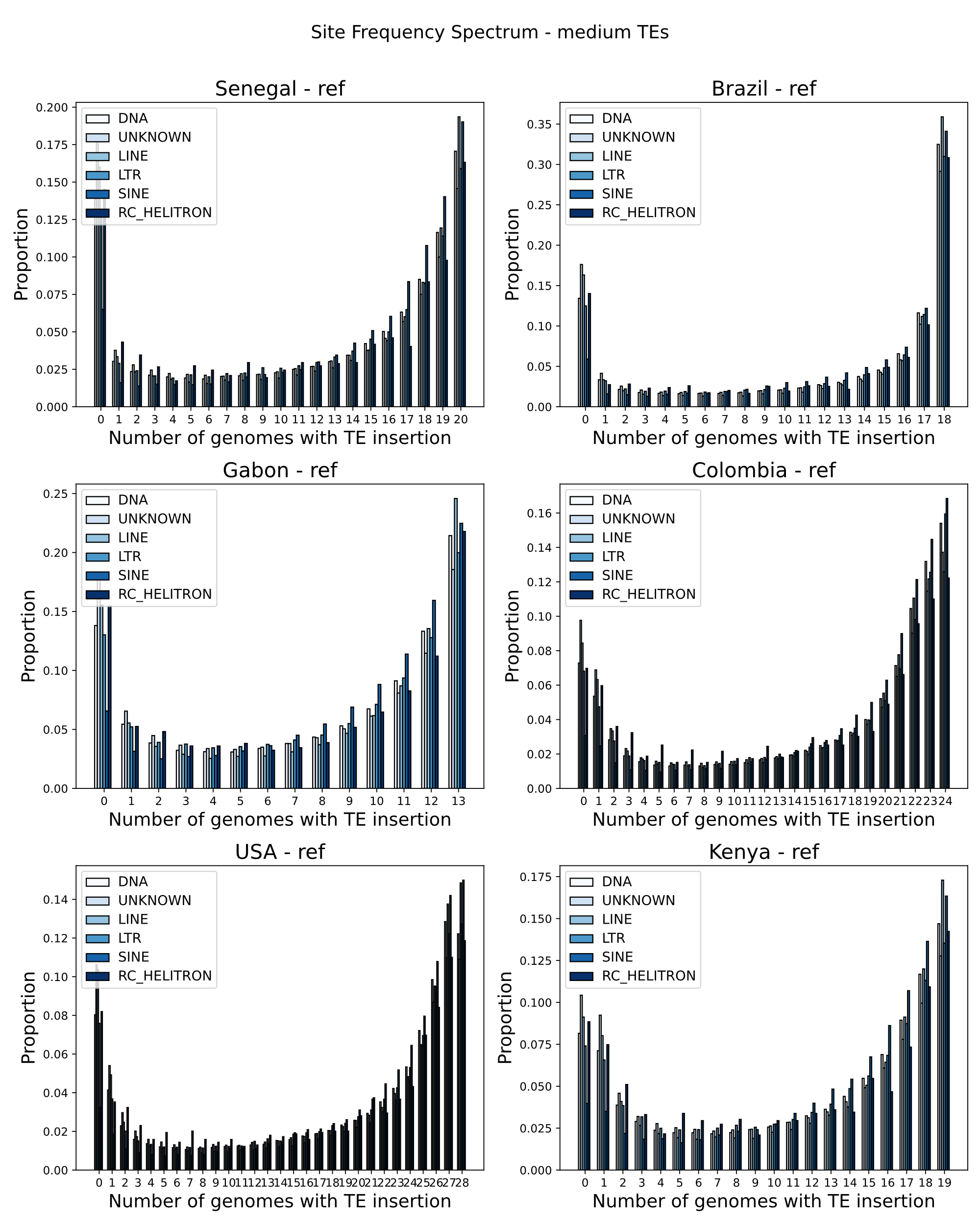

**Supplementary Figure 2:** Site Frequency spectrum of reference TEs based on the length. Between 200-600bp.

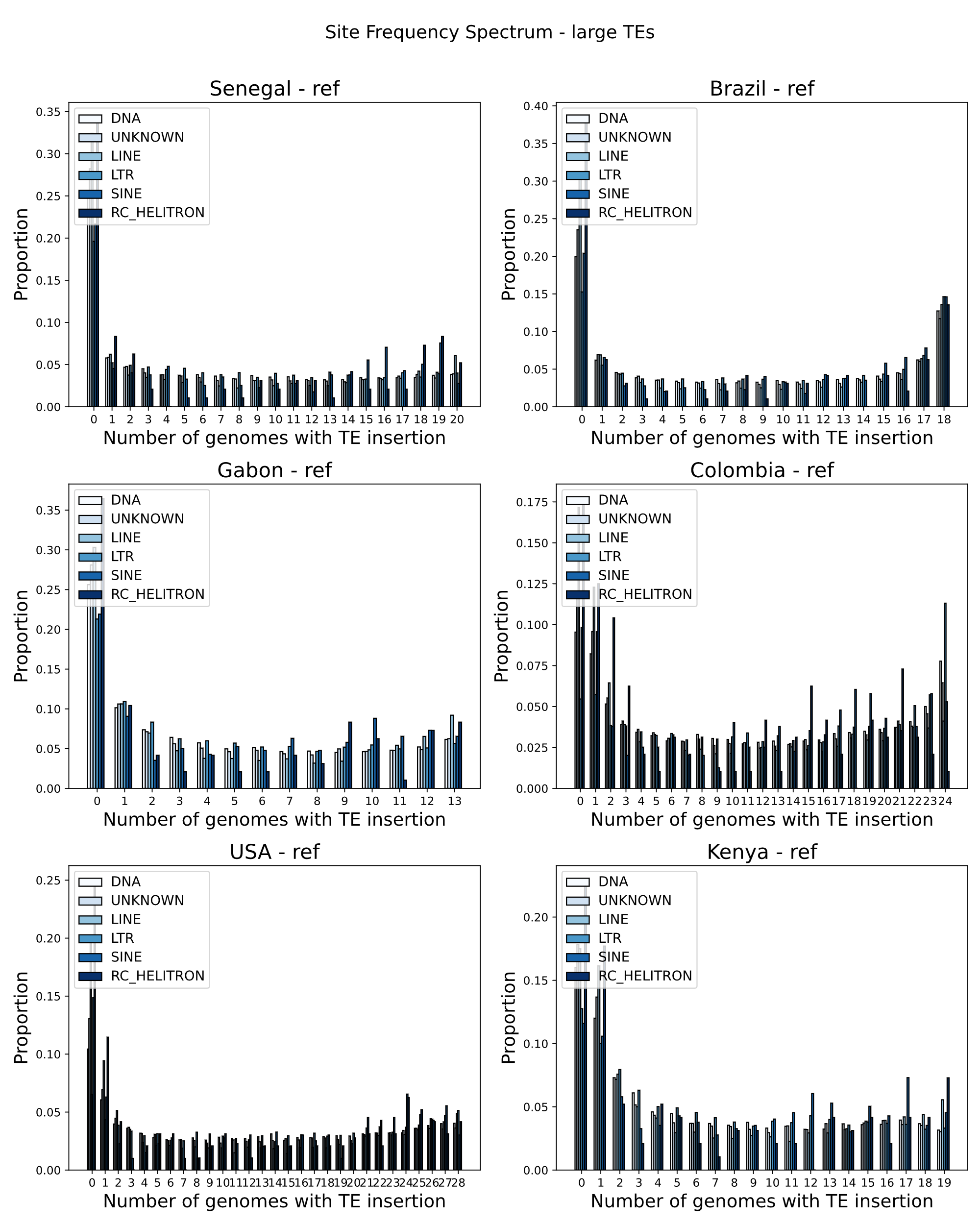

**Supplementary Figure 3:** Site Frequency spectrum of reference TEs based on the length. Bigger than 600bp.

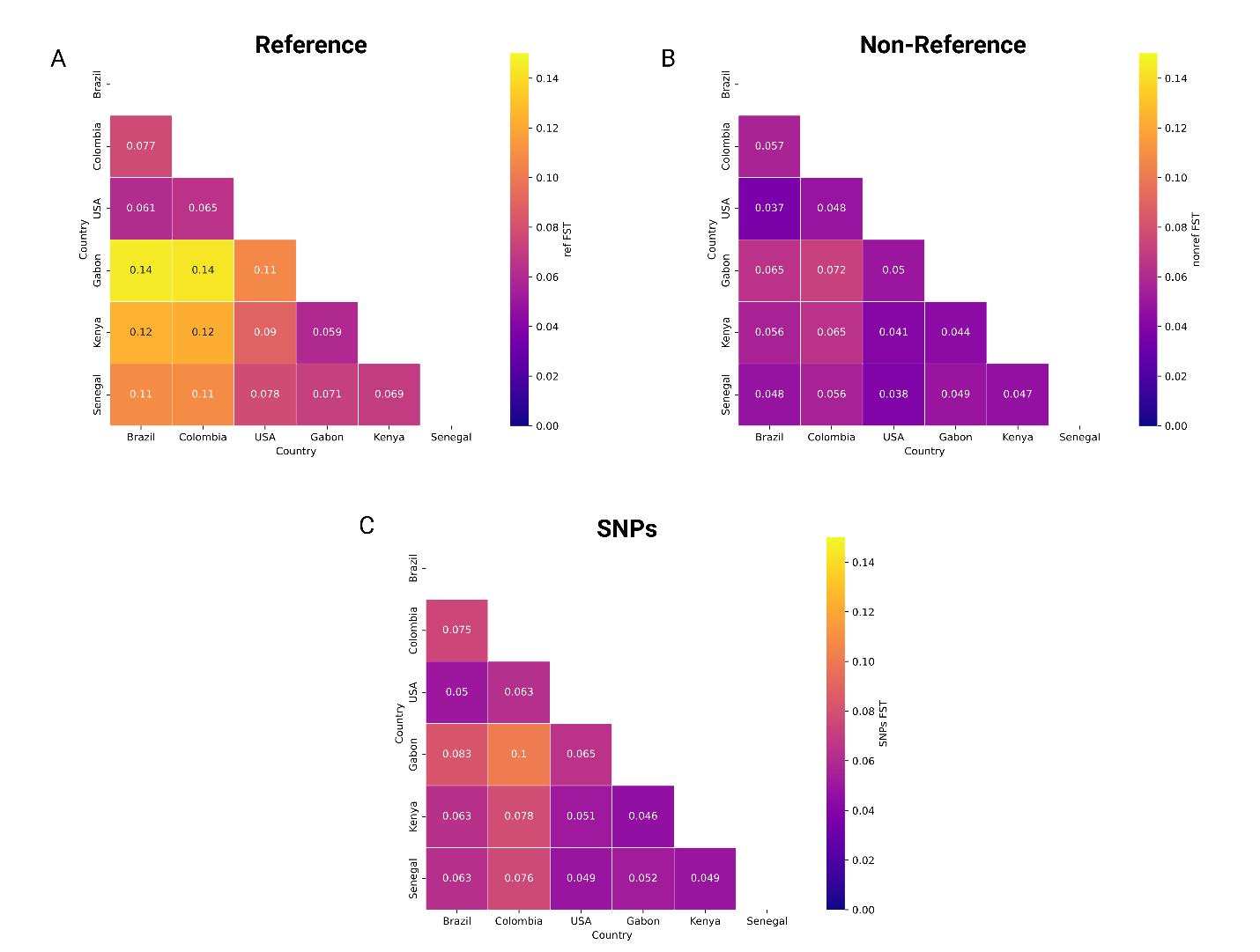

**Supplementary Figure 4:** Mean *F_ST_* values for a) reference TEs, B) non-reference TEs, and C) intergenic SNPs used to compare TEs to neutral variation.

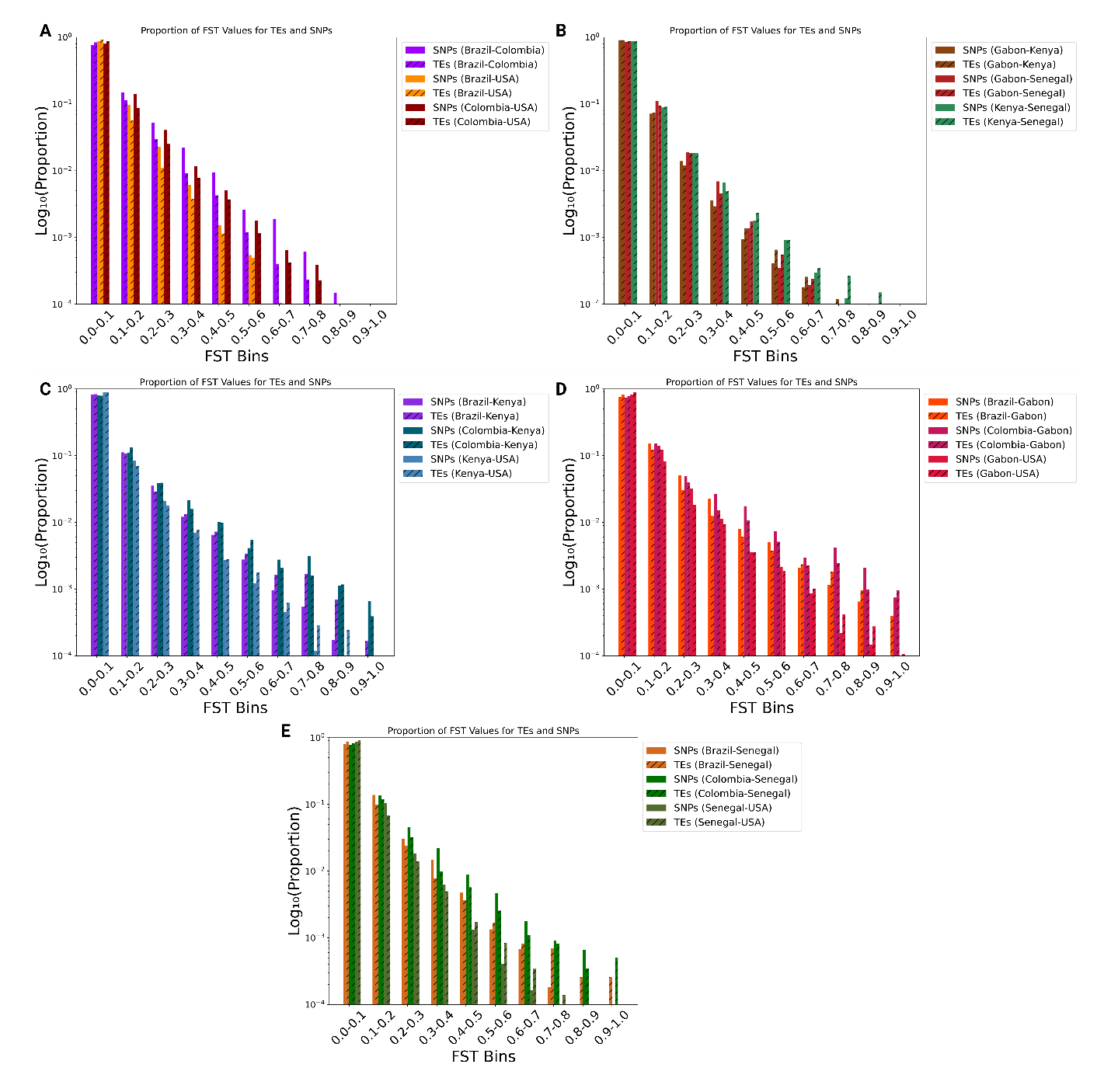

**Supplementary Figure 5:** Histogram of *F_ST_* values for combined reference and non-reference TEs (TEs), and neutral SNPs (SNPs). A) Comparisons among American countries. B) Comparisons among African countries. C) Comparisons between American countries and Kenya. D) Comparisons between American countries and Gabon. E) Comparisons between American countries and Senegal.

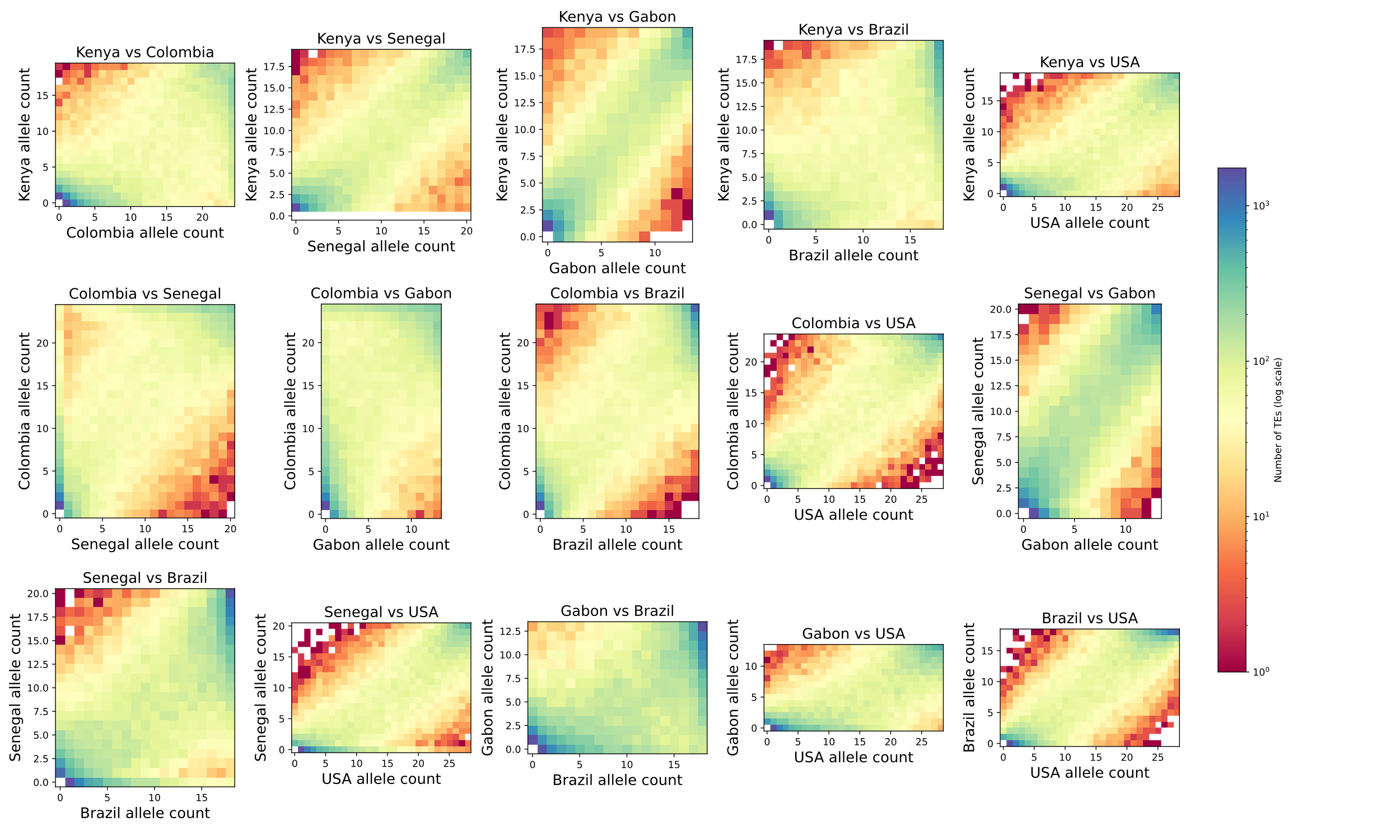

**Supplementary Figure 6:** Joint site frequency spectrum of reference transposable elements. All combinations of countries were analyzed. The coordinates of a given point in the plot specify a joint frequency, and the color at that point shows the number of TE insertions with that joint frequency.

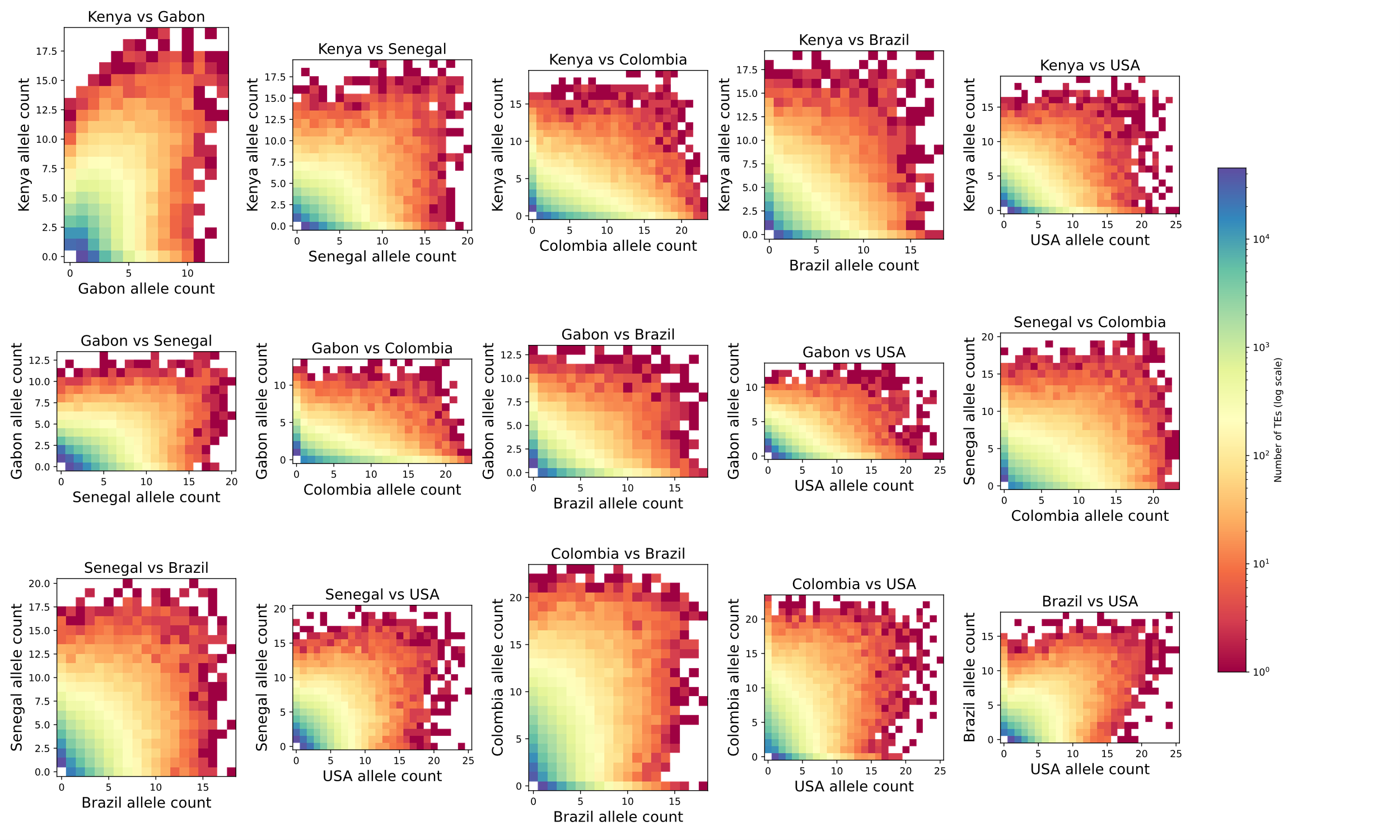

**Supplementary Figure 7:** Joint site frequency spectrum of non-reference transposable elements. All combinations of countries were analyzed. The coordinates of a given point in the plot specify a joint frequency, and the color at that point shows the number of TE insertions with that joint frequency.

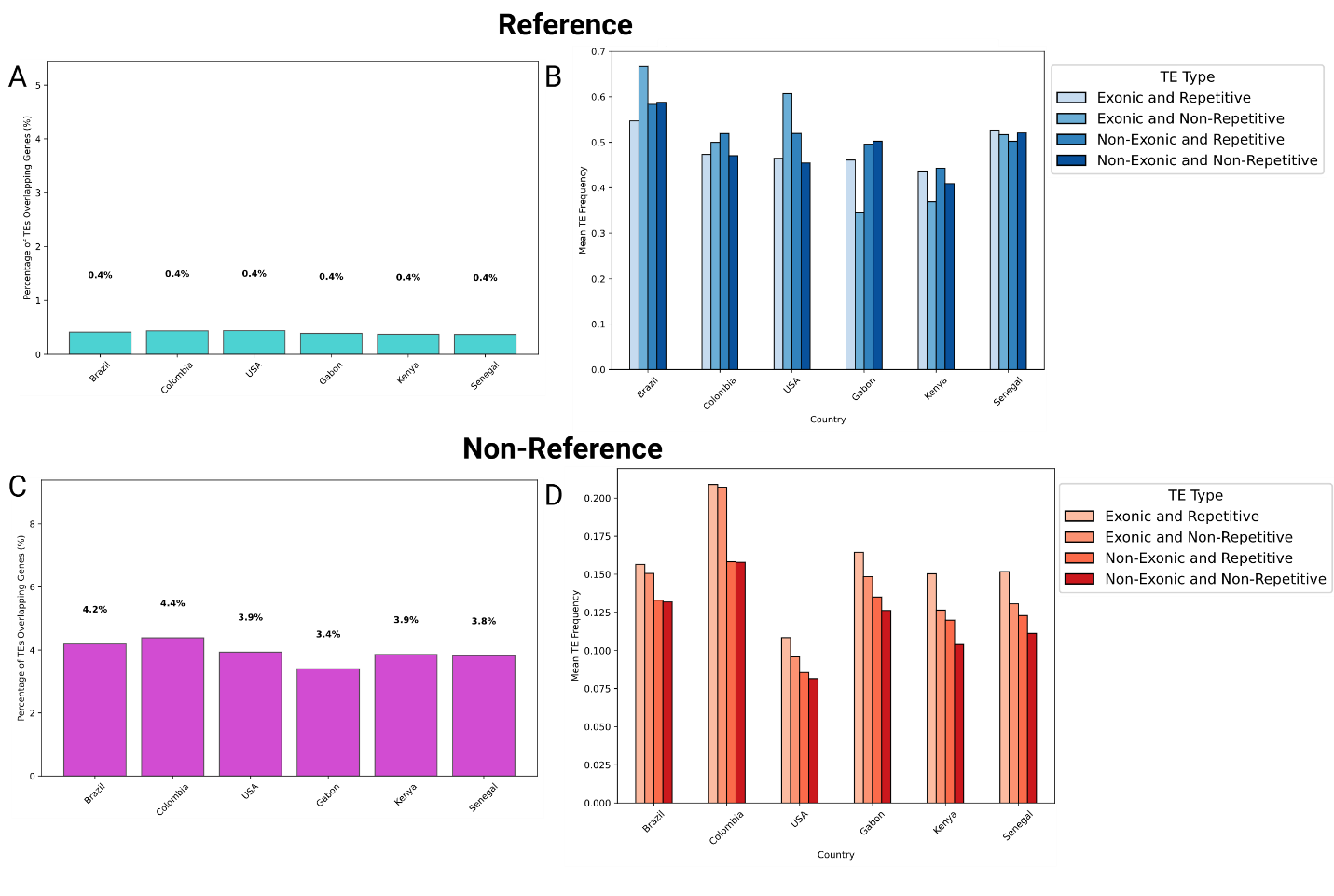

**Supplementary Figure 8:** TEs overlapping gene exon regions. A) Percentage of reference TEs for which the coordinates of the TE annotation at least partially overlapped one or more exons. B) Mean population frequency of reference TE insertions overlapping exons and annotated repetitive regions, overlapping exons and not overlapping repetitive regions, not overlapping exons and overlapping repetitive regions, and not overlapping exons or repetitive regions. C) Percentage of non-reference TEs whose breakpoint estimates overlap exons. D) Mean TE frequency of non-reference TEs in the same categories as in panel B.

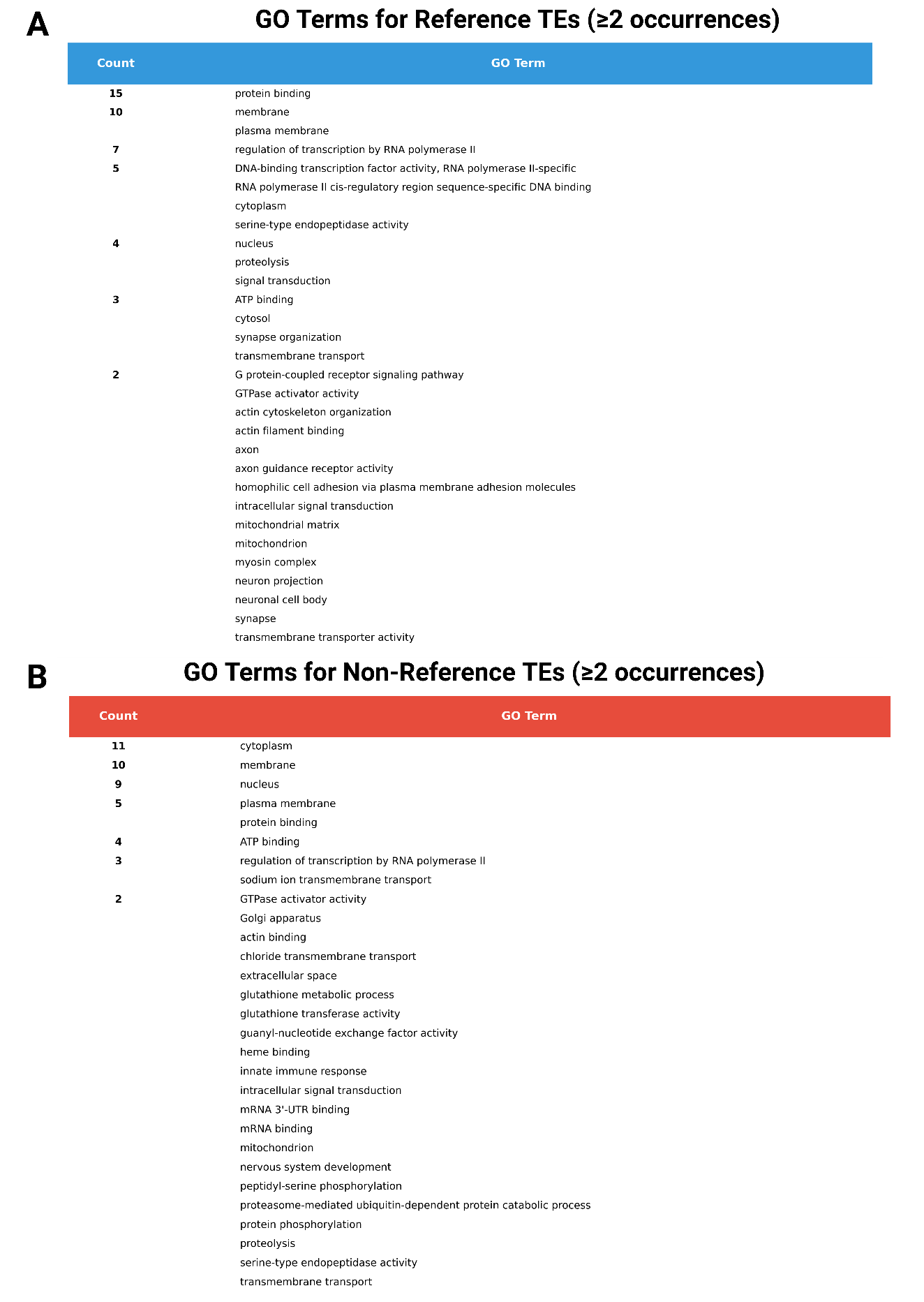

**Supplementary Figure 9:** GO terms from the genes that overlapped or were within 10 kb of at least one of the most differentiated TE insertion polymorphisms in our data set (combined across all country comparisons). A) Reference TEs. B) non-reference TEs.
